# *BiTSC*^2^: Bayesian inference of Tumor clonal Tree by joint analysis of Single-Cell SNV and CNA data

**DOI:** 10.1101/2020.11.30.380949

**Authors:** Ziwei Chen, Fuzhou Gong, Lin Wan, Liang Ma

**Affiliations:** NCMIS, Academy of Mathematics and Systems Science, Chinese Academy of Sciences, Beijing 100190, China; Institute of Zoology, Chinese Academy of Sciences, Beijing 100101, China; School of Mathematical Sciences, University of Chinese Academy of Sciences, Beijing 100049, China

## Abstract

The rapid development of single-cell DNA sequencing (scDNA-seq) technology has greatly enhanced the resolution of tumor cell profiling, providing an unprecedented perspective in characterizing intra-tumoral heterogeneity and understanding tumor progression and metastasis. However, prominent algorithms for constructing tumor phylogeny based on scDNA-seq data usually only take single nucleotide variations (SNVs) as markers, failing to consider the effect caused by copy number alterations (CNAs). Here, we propose *BiTSC*^2^, **B**ayesian **i**nference of **T**umor clonal **T**ree by joint analysis of **S**ingle-**C**ell **S**NV and **C**NA data. *BiTSC*^2^ takes raw reads from scDNA-seq as input, accounts for sequencing errors, models dropout rate and assigns single cells into subclones. By applying Markov Chain Monte Carlo (MCMC) sampling, *BiTSC*^2^ can simultaneously estimate the subclonal scCNA and scSNV genotype matrices, sub-clonal assignments and tumor subclonal evolutionary tree. In comparison with existing methods on synthetic and real tumor data, *BiTSC*^2^ shows high accuracy in genotype recovery and sub-clonal assignment. *BiTSC*^2^ also performs robustly in dealing with scDNA-seq data with low sequencing depth and variant dropout rate.

## 1 Introduction

The rapid development of single-cell DNA sequencing (scDNA-seq) technology has provided a refined perspective for unveiling the evolutionary mechanisms underlying cancer progression and for characterizing intra-tumor heterogeneity (1; 2). Although promising, the major single-cell whole genome amplification methods, e.g. DOP-PCR, MDA and MALBAC, still encounter various technical bottlenecks, which results in a high incidence of errors, such as dropout, false positive or false negative, in the sequenced single-cell DNA, and poses additional challenges for the downstream intra-tumor heterogeneity (ITH) inferences (3).

Early single-cell studies utilize information from single-cell SNV (scSNV) or single-cell CNA (scCNA) to infer tumor evolution with traditional phylogenetic methods (4; 5; 6; 7). In recent years, many computational methods proposed for inference of tumor evolutionary history from single-cell data have emerged. CHISEL (8) and SCICoNE (9) are the few methods that perform scCNA detection and also make inference to evolutionary histories. RobustClone(10), is a model free method which takes input of raw scSNV or scCNA genotype matrix to recover clone genotypes and infer tumor clone tree. BEAM is a Bayesian evolution-aware method on scSNV data, which improves the quality of single-cell sequences by using the intrinsic evolutionary information under a molecular phylogenetic framework (11). Many other methods based on scSNV data build maximum likelihood or Bayesian based models to account for sequencing noise as well as reconstruct tumor clone/cell tree. SCITE (12), OncoNEM (13), SCI*ϕ* (14) make infinite site assumption in their models, that is, mutation may only occur once at any locus and only binary genotypes are allowed in scSNV sites. SiFit (15) and SiCloneFit (16) constructed their models under the finite site assumption, which allows mutations to happen more than once at any locus.

These single-cell based methods can only take into account one source of information, either from scSNV or scCNA. In fact, these two types of markers all play important role in tumor generation, progression and metastasis, and they constitute crucial traits in characterizing tumor heterogeneity (17). Evolutionary inference with only one type of marker may lead to biased estimate. For example, suppose there is a true evolutionary process as shown in Figure 1A. The tumor tree 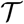 has five subclones, where subclone1 is a root node comprised of normal cells, and the others are cancerous subclones caused by point mutations and CNAs on three loci A, B and C. The SNV and CNA genotypes are shown in Figure 1B. The two SNVs occur on locus B and A which give rise to subclone2 and subclone4. If we infer the clone tree with only SNV data *Z*, we will generate a linear evolutionary history as in Figure 1C. This is biased as the copy loss at locus A and copy gain at locus C respectively give rise to two extra subclones (3 and 5). In this case, the full history can only be revealed by taking into account information from both SNV *Z* and CNA *L*.

**Figure 1:**
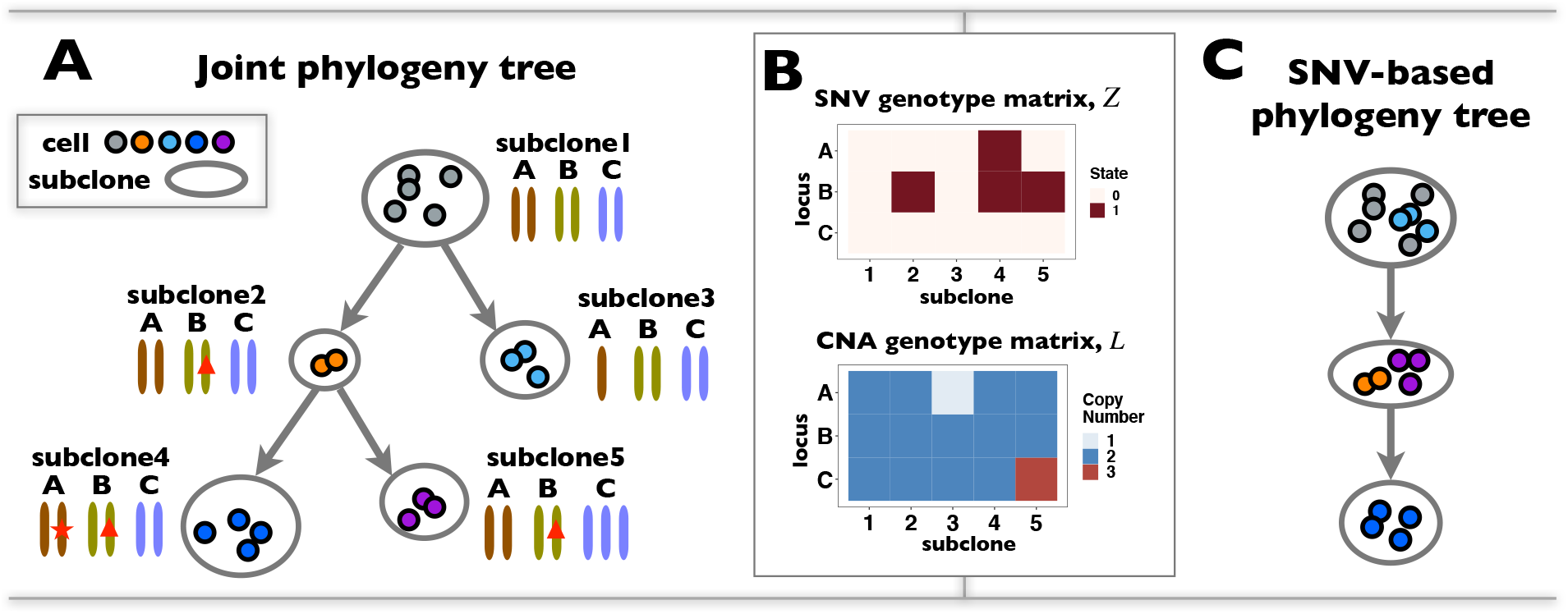
ScDNA-seq data display tumor heterogeneity. **(A)** Joint tumor phylogeny tree by SNV and CNA. **(B)** The SNV genotype matrix *Z* and the CNA genotype matrix *L*, where rows represent loci and columns are subclones. **(C)** The phylogeny tree generally obtained by SNV-based algorithms.

In fact, joint analysis of SNV and CNA in characterizing ITH is common with bulk sequencing. PyClone (18) applies Bayesian clustering to identify tumor clones/subclones based on SNVs and clonal CNAs (CNAs that are carried by all cancer cells). It provides insights to temporal ordering of mutations and subclones, but does not make inference to the tree structure. PhyloWGS (19) also employs a Bayesian framework with a tree structured stick breaking process as prior, which infers subclone cluster as well as the tree relationship of the subclones. Canopy (20) is a Markov Chain Monte Carlo (MCMC) algorithm for tumor evolution history inference, which accounts for both point mutation and raw copy number information. Recently, Zeng et al. (21) proposed a unified Bayesian feature allocation model on raw sequencing reads, SIFA. It provides a generating model that incorporates SNV and CNA to infer tumor phylogenetic tree.

To the best of our knowledge, the only method for tumor tree inference from scDNA-seq data that integrates SNV and CNA information is SCARLET (22). SCARLET optimizes for a loss-supported phylogeny. It first constructs a copy-number tree with existing methods and then refines such tree by resolving the multifurcations nodes using mutation profiles of the observed cells (22).

In this study, we propose **B**ayesian **i**nference of **T**umor clone **T**ree by joint analysis **S**ingle-**C**ell **S**NV and **C**NA, *BiTSC*^2^. It extends the SIFA model to Single-Cell DNA data, which integrates SNV and CNA information as well as accounts for sequencing error. *BiTSC*^2^ takes the observed total reads and mutant reads of single cells as input, models dropout rate and sequencing errors in scDNA-seq data and assigns single cells into subclones instead of deconvolution of mixed bulk samples. By applying MCMC sampling, *BiTSC*^2^ can simultaneously estimate the subclonal scCNA and scSNV genotype matrices, subclonal assignment and tumor subclonal evolutionary tree. In comparison with existing methods on synthetic and real tumor data, *BiTSC*^2^ shows high accuracy in genotype recovery and subclonal assignment. It is worth noting that *BiTSC*^2^ is also robust in dealing with scDNA-seq data with low sequencing depth and variant dropout rate.

## 2 Method

### 2.1 Overview of *BiTSC*^2^

In this section we give a brief introduction to *BiTSC*^2^, the general flowchart can be seen in Figure 2. *BiTSC*^2^ is a nonparametric Bayesian mixture model, which takes input of raw total and mutant read counts matrices *D_M×N_* and *X_M×N_* measured at *M* loci for *N* cells (Figure 2A). Due to the sharing of genetic information among homogeneous cells, we assume that there are *K* latent subclones in the *N* single cells drawn for sequencing (*K* ≪ *N*). The latent state of cell *n* is denoted by *C_n_* = *k*(*n* = 1,⋯, *N*). The parameter of the Categorical distribution denoted by *ϕ* represents the prevalence of each subclones in the sample, and is given a symmetric Dirichlet prior with hyper-parameter *γ*. Each subclone consists of a group of cells with identical genotype, and distinct subclones differ in SNV or CNA markers on at least one of the *M* measured loci. We employ a tree coupled generating model to generate subclone genotypes *Z* and *L*, which are jointly modeled by *Z^o^* and *L^o^*, the SNV and CNA origin matrices, and the unknown clone tree 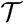. In addition, as we are modeling single-cell sequencing data, our model also accounts for sequencing error rate (*ε*) and dropout rate (*ρ*) (Figure 2B).

**Figure 2:**
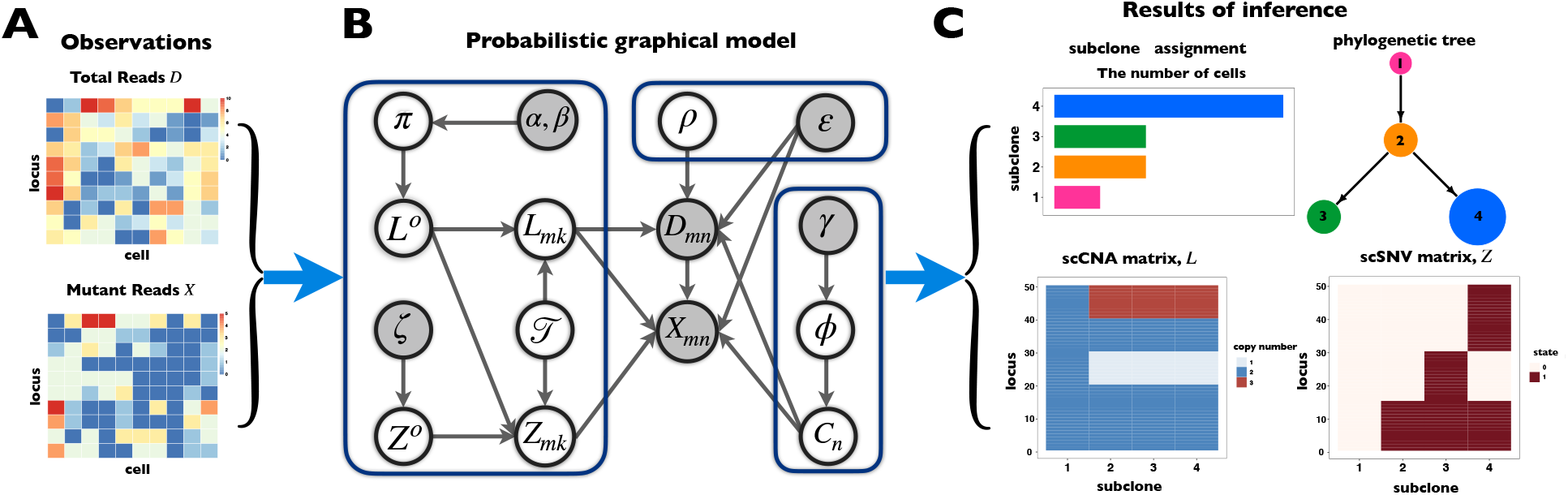
Overview of the computational framework of *BiTSC*^2^ that identifies subclones, recover subclonal genotypes of CNA and SNV, as well as reconstructs subclonal evolutionary trees using tumor scDNA-seq read count data. **(A)** The input of the algorithm, total reads matrix *D* and mutant reads matrix *X*. **(B)** The probabilistic graphical model shows the dependency among parameters, where the shades nodes stands for observed or fixed values, the unshaded nodes represent the latent parameters. **(C)** The inference output of the algorithm, containing dropout rate, subclone assignment, subclonal phylogenetic tree and genotype matrix of CNA, *L*, and SNV, *Z*.

The ultimate goal of *BiTSC*^2^ is to infer the subclone prevalence *ϕ*, the subclone assignment of cells *C*, the SNV and CNA genotypes of subclones *Z* and *L*, the subclone tree 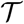 and also the dropout rate *ρ* (Figure 2C). By assigning priors to *Z^o^, L^o^*, 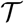, *C* and *ρ*, and given a sequencing error rate *ε*, these can be estimated from a posterior distribution *p*(*ϕ, C, L, Z*, 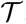, *ρ|D, X, ε*), which corresponds to *p*(*ϕ, C, L^o^, Z^o^*, 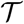, *π, ρ|D, X, ε*).

### 2.2 Tree coupled generating model of genotypes

The subclone genotypes *Z* and *L* are generated according to the SNV and CNA origin matrices *Z^o^* and *L^o^* as well as the clone tree 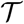. By assuming a total of *K* subclones in the tree, 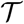 is represented by a length-*K* vector, where 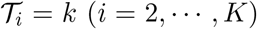 indicates the parent of subclone *i* is *k*. We fix subclone1 to normal cell and place it at the root of the tree 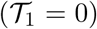. We give a uniform prior to all possible trees with *K* nodes.

We assume SNV and CNA mutations arise independently. Each mutation (including SNV and CNA) originates only once in a specific subclone besides normal cell. The mutation will be inherited by all descendant subclones of its origination. Then we represent the originations of SNV and CNA changes at locus *m* with 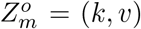 and 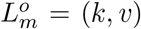, where 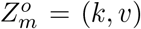 indicates mutation at locus *m* occurs from subclone *k* and gains *v* mutant copies, and 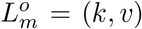 indicates the CNA arises in subclone *k* and gains (or loses) *v* normal copies.

For SNV state, we take the prior of *Z^o^* as 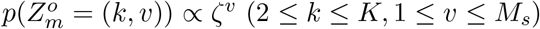, where *ζ* is the somatic point mutation rate and is predetermined in (0,1), *M_s_* represents the maximum number of possible mutant copies (21). We further assume that obtaining multiple copies of the mutant alleles is a less likely event, therefore we set the probability of obtaining *v* copies of the mutant alleles to *ζ^v^*. The specification of the mutation probability is independent of *k*, which makes it equally likely for the SNV to originate from any subclones (except for the normal).

Similarly, for CNA state, we set *M_c_* as the maximum total alleles. Since CNA occurs on chromosome fragments, we can use the information to better infer the status of CNA. We sort the loci in the order of chromosomal positions, and divide the genome into *S* segments, that is, {Δ_1_, Δ_2_, ⋯, Δ_*S*_}. If the loci *i* and *j* are located on the same segment, we assume that they share the same CNA status. There exist many other methods that can be applied to estimate the segment information, such as, HMMcopy (23; 24). Let (0,0) represent no CNA event. For each genome segment, Δ_*s*_, we assign the copy number status prior as 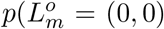 for all *m* ∈ Δ_*s*_) = *π*, and uniform on the other possible combinations of *k* and *v*. The hyper-parameter *π* is given a prior of Beta(*α, β*).

The independent origination of SNV and CNA at *M* loci coupled with the *K* nodes clone tree 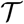 will derive the *M* × *K* genotype matrices *Z* and *L*. We select the optimal number of subclones *K* according to a modified Bayesian Information Criterion (BIC, see Supplementary Note 4 for details).

### 2.3 Zero-inflated modeling of single-cell sequencing reads

We will next introduce the likelihood of observing *D_mn_* total reads and *X_mn_* mutant reads at locus *m* of cell *n*, given the genotypes of subclones.

By given the latent subclone state *C_n_*, e.g. *C_n_* = *k*, the total reads *D_mn_* should be positively correlated with copy number *L_mC_n__* and the cell specific average coverage *ψ_n_* (which should be given a priori) of cell *n* (21; 25; 26). We apply Poisson distribution to model the total read as:

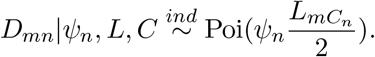

Note that when the total copy number of a single cell is equal to 2, the mean and the variance of the Poisson distribution are equal to *ψ_n_*.

We then denote the expectation of mutant allele at locus *m* for cell *n* as *p_mn_* = *Z_mC_n__*/*L_mC_n__*. The likelihood of observing *X_mn_* reads is thus modeled by Binomial distribution as:

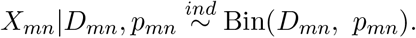

Since in single cell sequencing, data are often disturbed by high noise due to events such as dropout or sequencing error, especially at low sequencing depth. We adopt a zero-inflated distribution, which adds an additional parameter *ρ* to the existing Poisson distribution to control the amount of excessive zeros, named zero-inflated Poisson (ZIP) distribution (27) to model dropout rate. In addition we account for sequencing error *ε* in scDNA data, which may cause false positive reads. The ZIP likelihood of *D_mn_* can be defined as follows:

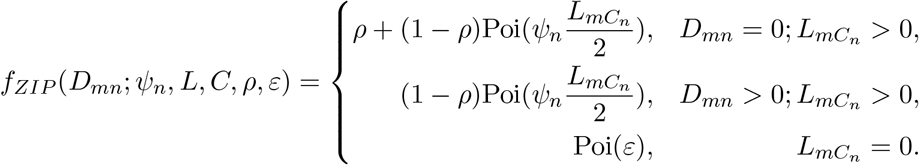

We also account for sequencing error in modeling mutant read. To do so, we assume that if mutation m is absent in cell *n*, i.e., *p_mn_* = 0, the probability of observing a variant read corresponds to the per-nucleotide rate of sequencing error *ε*. If mutation *m* presents in cell *n* and 0 < *p_mn_* < 1, we model the variant counts at locus *m* with a corrected Binomial distribution, where the underlying reads frequency of *p_mn_* is corrected by sequencing errors producing any of the other two bases. And if mutation *m* presents with *p_mn_* = 1, that is, *Z_mC_n__* = *L_mC_n__*, we give a small probability *ε* to observe normal reads. We thus write the likelihood for observing *x_mn_* as:

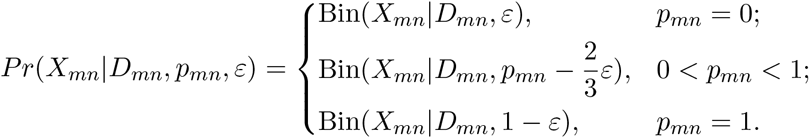

### 2.4 Inference

We apply the MCMC procedure to estimate the unknown parameters in *BiTSC*^2^. The posterior of the unknowns is sampled with differed strategies.

For genotype origin matrices *Z^o^* and *L^o^*, we apply Gibbs sampler, which updates one locus at a time. If the CNAs are in a segmented form, then at each step we will update all loci within the same segment instead. The hyper-parameter *π* of *L^o^* is also sampled by Gibbs.

For dropout rate *ρ*, since it is difficult to sample from its full conditional distribution, we adopt Metropolis sampling step with a uniform proposal of *ρ* in the interval [0,1]. We apply a mixed sampling strategy for 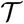 as in (21), where the tree is updated by randomly applies a Metropolis-Hastings sampler or a slice sampler.

In sampling of the subclone prevalence, instead of updating the entire vector *ϕ* at once, we sample additional Gamma parameters *θ_k_*~*Gamma*(*γ*, 1), *k* = 1⋯*K*, one at a time. And let 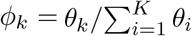. This move is equivalent to sampling *ϕ* with prior *Symmetric – Dirichlet*(*K*; *γ*), and often leads to better mixing of the MCMC (21). Each *θ_k_* is update by Metropolis-Hastings sampling with a Gamma proposal and an adaptive step size. Each element *C_n_*(*n* ∈ {1,2, ⋯, *N*}) of *C* is taken from the Categorical distribution with parameter *ϕ*. We employ Gibbs sampling to update each *C_n_* one by one (the detail sampling process for all parameters can refer to Supplementary Note 1).

In order to avoid sampled states being trapped at some local optimum in MCMC, we adopt parallel tempering technique, which run multiple chains with different temperatures, and exchange samples between them (21; 28). We use heuristic initialization for each parallel chain before MCMC sampling (Supplementary Note 2). The derivation of the fully conditional distribution for all model parameters can refer to Supplementary Note 3. And the optimal number of subclones *K* is selected by performing a modified Bayesian Information Criterion (BIC) (Supplementary note 4). We use the posterior mode for 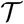 and *C*, and the posterior median for *Z* and *L*, as the final estimates. The software is available at https://github.com/ucasdp/BITSC2.

### 2.5 Benchmark *BiTSC*^2^

#### 2.5.1 Simulation Data

In order to systematically evaluate the performance of *BiTSC*^2^, we simulated 150 datasets with variant number of cells (*n*), sequencing depths (Ψ), dropout rate (*ρ*), as well as the number of loci (*m*) and the number of subclones (*K*). The 150 datasets are divided into five groups (denoted G1-G5), each of which contains 30 datasets. In each group we change one parameter and keep other parameters fixed. In addition to the variable parameter, we set the default parameters in each group as follows: number of cells (*n*) is 100, number of loci (*m*) is 100, dropout rate (*ρ*) is 20%, sequencing depths of all single cells (Ψ) are 3, and the number of subclones (*K*) is 4. The ground truth (including *Z, L*, and tree structure) of G1-G3 is shown in Figure S1, and the ground truth of G4 and G5 is shown in Figure S2 and S3, respectively. Specific parameter settings of G1-G5 can refer to Table S1.

We also simulated another 10 datasets, denoted as G6, to test the ability of *BiTSC*^2^ to identify the subclones which only contain CNA allocations. Figure S4 shows the ground truth of the datasets, which contains the phylogenetic tree and subclonal genotype matrices of CNA and SNV.

#### 2.5.2 Real Data

We apply *BiTSC*^2^ on the scDNA-seq data from metastatic colorectal cancer patient CRC2 in (29). This data include 141 cells from the primary colorectal tumor and 45 cells from a matched liver metastasis by single cell DNA target sequencing of 1,000 caner genes with an average sequencing depth of 137×.

### 2.6 Evaluations

We compare the performance of our algorithm to RobustClone, BEAM (10; 11), which are two algorithms that perform very well in genotype recovery problem in systematic comparison of RobustClone, under our different simulated scenarios. The evaluations include: 1) adjusted Rand index (ARI) (30; 31) to measure the similarity of subclone assignment between ground truth and estimation (details can refer to Supplementary Note 5); 2) the error rate of the recovered scSNV genotype matrix to calculate the accuracy of scSNV genotype recovery.

## 3 Results

### 3.1 *BiTSC*^2^ recovers genotype matrix and assigns cells with high accuracy on synthetic datasets

We compare *BiTSC*^2^ to RobustClone and BEAM, two most recent methods on tumor evolutionary inference with single-cell data, on 6 groups of synthetic data. We give *BiTSC*^2^ the real segmentation information according to the CNA genotype matrices (Figure S1-S4). The prior settings and the MCMC configurations of *BiTSC*^2^ for simulation data are shown in Table S2-S3. We perform *BiTSC*^2^ with the number of subclones *K* in range from 3 to 7, and select the best fitted *K* by BIC.

Figure 3 shows the results of comparisons in G1-G5, with top rows of Figure 3A and B showing the overall performance, and bottom rows display the detailed benchmarks at differed settings. In general, compared with the other two algorithms, *BiTSC*^2^ has the high robustness in subclone assignments (Figure 3A) and accuracy in recovering SNV genotypes (Figure 3B).

**Figure 3:**
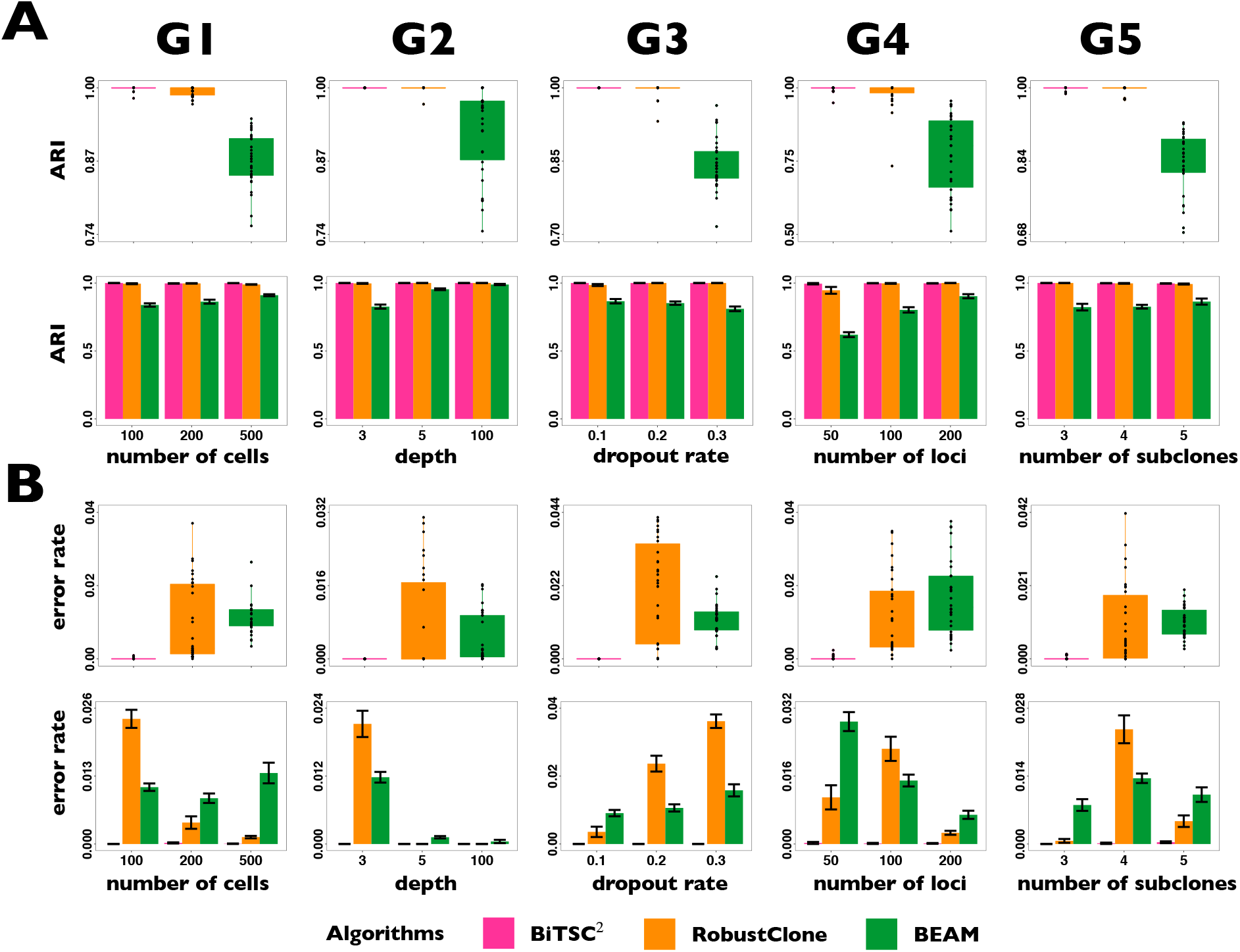
Comparison of performs on G1-G5 for subclone assignment and scSNV genotype recovery among *BiTSC*^2^ with real segment information input, RobustClone and BEAM. **(A)** The boxplot and barplot of three algorithms for ARI of subclone assignment. **(B)** The boxplot and barplot of three algorithms for error rate of recovered scSNV genotype matrix.

Specifically, for subclone assignment, *BiTSC*^2^ and RobustClone show high robustness and consistency with ground truth. BEAM is slightly less consistent with ground truth in subclone assignment than *BiTSC*^2^ and RobustClone, but gets improved with the increase of number of cells (*n*), number of loci (*m*) and sequencing depths (Ψ) (second row in Figure 3A). For the accuracy of genotype recovery, *BiTSC*^2^ shows much higher accuracy and robustness than RobustClone and BEAM. The accuracies of RobustClone and BEAM improve with the increase of number of loci (*m*) and sequencing depths (Ψ), but reduce with the increase of number of subclones (*K*) and dropout rate (*ρ*). The difference is that accuracy of RobustClone gets much better with the increase of number of cells (*n*), while BEAM shows a large variance when the number of cell (*n*) is 500, and the average error rate is higher when the number of cell (*n*) are 100 and 200 (second row in Figure 3B).

Figure 4 shows the comparison result of G6, which includes 10 replicates with a CNA induced lineage (Figure S4). *BiTSC*^2^ is able to correctly detect subclone which caused by only CNA mutation, and can accurately assign cells into four subclones and recover SNV and CNA subclonal genotypes. While RobustClone and BEAM can only analyze one source of data (SNV), although their performance in recovery of SNV genotypes were acceptable, their performance in subclone assignment were unsatisfactory (Figure 4AB).

**Figure 4:**
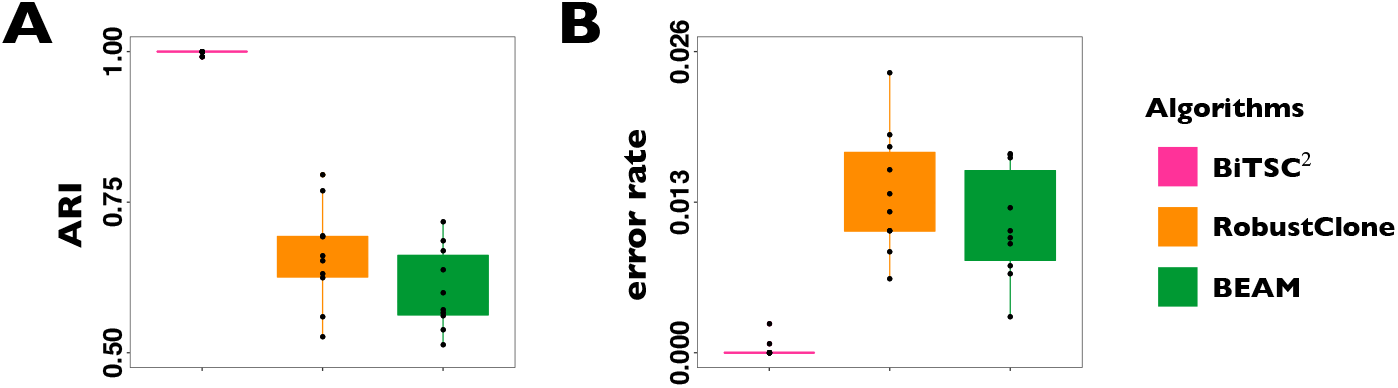
Comparison of performs on G6 for subclone assignment and scSNV genotype recovery among *BiTSC*^2^ with real segment information input, RobustClone and BEAM. **(A)** The boxplot of three algorithms for ARI of subclone assignment. **(B)** The boxplot of three algorithms for error rate of recovered scSNV genotype matrix.

In the above comparisons *BiTSC*^2^ was given the real segmentation information as input. However, this information may not always be reliably estimated. In that case, we could take the more refined raw bins (the bins after binning step before segmentation and CNA calling) as segments or use locus specific segments (each gene/SNV locus as a segment). Here we additionally evaluate *BiTSC*^2^ on G1-G6, with a locus specific segment setting, that is, instead of updating loci in the same segment, we update one locus in *L^o^* at a time. The results show that *BiTSC*^2^ maintains high accuracy robustness (Figure S5, S6). For genotype recovery, the overall performance of *BiTSC*^2^ is higher than RobustClone and BEAM, but with sequencing depth 100 and the number of subclones 3, there is decline in inference accuracy, where the performance of *BiTSC*^2^ fall slightly behind as compared to RobustClone and BEAM. This might due to the setting of prior distribution *π* (the prior probability of a segment with a copy neutral state) is too close to 1, which aims to control the false positives of CNA calls. When the multiple loci in a segment with CNA events are jointly updated, the likelihood of the model will be greatly improved, where the same CNA state will be assigned to all the locus in same segment. When a single locus does not contribute much to the likelihood of the model, it is likely that the CNA will not be allocated. Therefore, the performance of *BiTSC*^2^ reduced slightly for locus specific segment setting as compared to cases where correct segment information can be provided.

### 3.2 *BiTSC*^2^ recovers single cell phylogeny of metastatic colorectal cancer

We apply *BiTSC*^2^ to analyze the scDNA-seq data of colorectal cancer patient CRC2 in (29) with primary and metastatic samples. After filtering for some low-coverage data, the sequencing data of 182 single cells and 36 SNV loci were retained for further analysis. We apply *BiTSC*^2^ directly to the raw reads covering these loci and apply the locus specific segment setting to CNAs. The cell specific sequencing depth of each single cells can be found in the Supplementary Table S4 in (29). The *BiTSC*^2^ is performed with prior and MCMC settings shown in Table S4-S5.

*BiTSC*^2^ reconstructs a clone tree of 9 subclones as shown in Figure 5A (see Figure S7 for the BIC values). Figure 5B displays the prevalence of cells in each subclone. The metastatic aneuploid cells are mainly distributed in subclones 8 and 9, while the primary aneuploid cells are predominantly clustered in subclone6 (Figure 5C). Although the cells occupied the other subclones were labeled diploid by (29), we still find some CNA events at many targeted genes (Figure 5D). Extensive point mutations were identified in primary (subclone6) and metastatic (subclones 8 and 9) tumor cells (Figure 5E).

**Figure 5:**
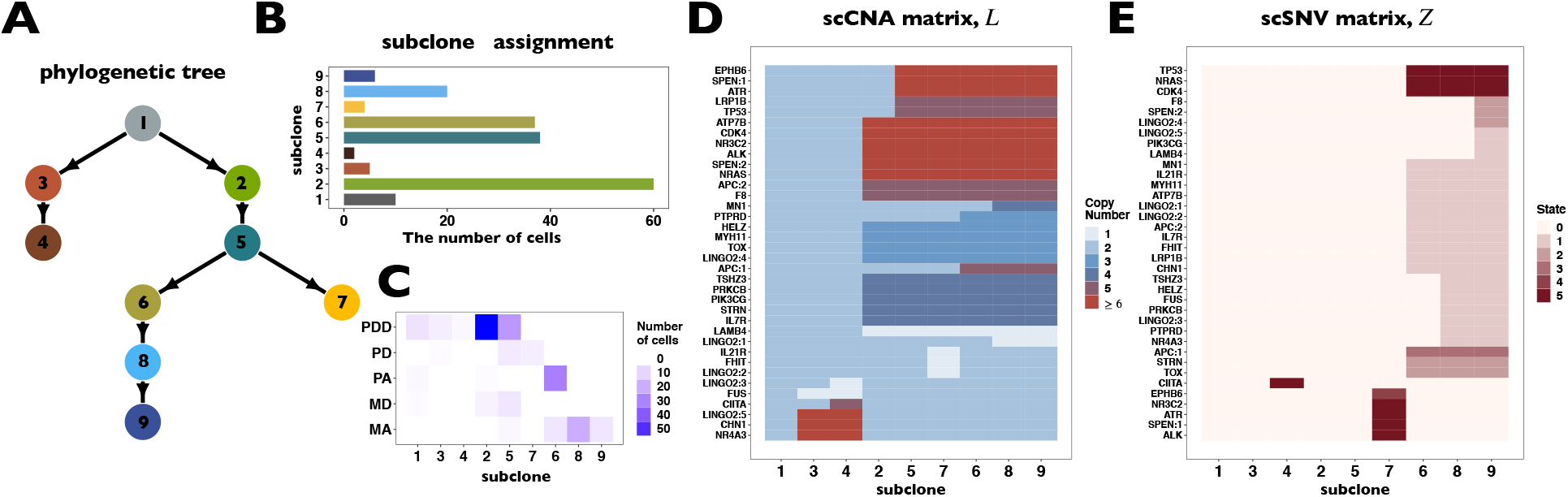
*BiTSC*^2^ reconstructs tumor phylogeny of metastatic colorectal cancer. **(A)** The phylogeny tree of metastatic colorectal cancer reconstructed by *BiTSC*^2^. **(B)** The subclone assignment. **(C)** The number of overlapped cells contained in subclones identified by *BiTSC*^2^ and cells contained in the targeted region, where PD stands for Primary Diploid, PA stands for Primary Aneuploid, MD stands for Metastatic Diploid, and MD stands for Metastatic Aneuploid in (29). **(D)** The CNA subclonal genotype matrix calculated by *BiTSC*^2^. **(E)** The SNV subclonal genotype matrix calculated by *BiTSC*^2^.

Interestingly, our inferred tumor clone tree and genotypes show that metastatic cells (subclones 8 and 9) majorly share the same CNA events on LINGO2:1 and MN1, which arise in the descendant of primary sites (subclone6). Contrary to the polyclonal seeding (that is, two independent clones with different mutations migrate from primary colon cancer to liver metastases at different time points) conclusion based on SCITE tree in the original study (29), our result indicates that the liver metastasis from colon was a single event, which supports the monoclonal seeding hypothesis and is consistent with the inference based on the SCARLET tree (Figure S8) (22).

Besides the metastatic lineage, we also identified two other lineages with unique mutations respectively. One lineage leads to subclone4 which consists of a small proportion of cells that carries SNV on CIITA. Such lineage was also identified by SCITE and SCARLET trees (Figure S8AB). The other lineage, subclone7, descended from subclone5 and evolved in parallel to the primary tumor lineage (Figure 5A). Subclone7 can be identified by point mutations on ALK, ATR, EPHB6, NR3C2, and SPEN:1 and share no common mutations with major primary or metastatic aneuploid tumor cells. This result was also supported by SCITE tree which constructed with only scSNV data. Although a similar lineage was seen in SCARLET tree, however, it also possesses many extra shared mutations to primary tumor cells, such as APC:2, NRAS, or TP53 (Figure S8B).

## 4 Discussion

Computational method based scDNAseq data for tumor ITH and evolutionary history inference can provide important insights to the understanding of tumor progression and metastasis mechanism, and provide guidance to tumor treatment and response. Most of such methods only utilize one source of information, either SNV or CNA, which may lead to biased estimation of the true evolution history of cancer. In this study, we propose *BiTSC*^2^, a Bayesian-based method that integrates SNV and CNA markers from scDNA-seq data to jointly infer tumor clone tree. *BiTSC*^2^ is a unified Bayesian framework, which takes the raw total reads and mutation reads generated by sequencing as input and takes into account sequencing errors and models dropout rate. It also optimizes SNV and CNA subclonal genotypes matrices, assigns cells to subclones, and constructs subclonal tree. *BiTSC*^2^ has a high accuracy for subclone assignment and SNV subclonal genotypes matrix recovery compared to existing methods such as RobustClone and BEAM. *BiTSC*^2^ can handle low-depth single-cell sequencing data with high performance. *BiTSC*^2^ also provides high accurate and robust estimation of the dropout rate in scDNA-seq data (Figure S9).

When *BiTSC*^2^ updates the CNA subclone genotype matrix *L*, it prefers to update all the loci in the same segment together, because the loci in the same segment share the same CNA status. When we have no information about the real segment information, there are many existing methods can be applied to perform segmentation, for example, HMMcopy, copynumber, etc. (24). In cases when segment information can not be reliably obtained, *BiTSC*^2^ can also update *L^o^* and *L* locus by locus in the same way as updating *Z^o^* and *Z*. In the results on synthetic data, we show that the accuracy and robustness of updating one locus at a time are still higher than RobustClone and BEAM in most cases (see Section 3.1, Figure S5, S6). In this way, *BiTSC*^2^ may provide a raw estimate of CNA segments based on the inferred CNA genotype matrix *L*.

We have applied *BiTSC*^2^ to a real scDNA-seq of a colorectal cancer patient. This data was originally analysed in (29) tumor clone tree constructed by SCITE (12) based on scSNVs. They found two distinct branches lead to metastatic cells, and consider this as the evidence of polyclonal seeding events. SiCloneFit (16) re-analyzed the same data (scSNVs) using a finite site model and also confirmed polyclonal seeding of metastatic tumors. Our result, however, indicates a monoclonal seeding of metastasis tumor cell. Such result was supported by SCARLET (22), which also integrates both SNV and CNA information. In this comparison, we show the importance of joint modeling and integrating of both SNV and CNA markers.

## 5 Acknowledgements

This work was supported by the National Key R&D Program of China under Grant 2019YFA0709501, 2018YFB0704304, NSFC grants (Nos. 81673833, 11971459, and 12071466), NCMIS of CAS, LSC of CAS, and the Youth Innovation Promotion Association of CAS.

