## supplemental material for "*BiTSC*^2^: Bayesian inference of Tumor clonal Tree by joint analysis of Single-Cell SNV and CNA data"

#### Supplementary Information

Ziwei Chen[1,3] Fuzhou Gong[1,3] Lin Wan[1,3*]
Liang Ma[2 *]

[1] NCMIS, Academy of Mathematics and Systems Science, Chinese Academy of Sciences, Beijing 100190, China

[2]Institute of Zoology, Chinese Academy of Sciences, Beijing 100101, China

[3]School of Mathematical Sciences, University of Chinese Academy of Sciences, Beijing 100049, China

### Contents

Supplementary Methods ... 3

Supplementary Note 1: Inference of parameters ... 3

Sampling of subclone assignment $C$ ... 3

Sampling of SNV and CNA origin matrices $L^o$, $Z^o$ ... 3

Sampling of clone tree $\mathcal{T}$ ... 4

Sampling of dropout rate $\rho$ ... 4

Supplementary Note 2: Heuristic initialization process for MCMC parallel chains ... 5

Supplementary Note 3: Derivation of the fully conditional distribution for all model parameters ... 5

Supplementary Note 4: Model selection ... 6

Supplementary Note 5: Evaluations of ARI ... 7

Supplementary Figures ... 8

Figure S1: The ground truth of simulation datasets in G1-G3, containing subclonal phylogenetic tree and genotype matrix of CNA $L$ and SNV $Z$. ... 8

Figure S2: The ground truth of simulation datasets in G4, containing subclonal phylogenetic tree and genotype matrix of CNA $L$ and SNV $Z$. ... 9

Figure S3: The ground truth of simulation datasets in G5, containing subclonal phylogenetic tree and genotype matrix of CNA $L$ and SNV $Z$. ... 10

Figure S4: The ground truth of simulation datasets in G6, containing subclonal phylogenetic tree and genotype matrix of CNA $L$ and SNV $Z$. ... 11

Figure S5: Comparison of performs on G1-G5 for subclone assignment and scSNV genotype recovery among $BiTSC^2$ with updating one locus at a time, RobustClone and BEAM. ... 12

Figure S6: Comparison of performs on G6 for subclone assignment and scSNV genotype recovery among $BiTSC^2$ with updating one locus at a time, RobustClone and BEAM. ... 13

Figure S7: The BIC of metastatic colorectal cancer data calculated in Model selection step. ... 14

Figure S8: The phylogeny tree inferred by SCITE and SCARLET on the dataset from a Metastatic Colorectal Cancer Patient. ... 15

Figure S9: The dropout rate $\rho$ estimated by $BiTSC^2$ on the datasets from G3 groups. ... 16

Supplementary Tables ... 17

Table S1: The setting of simulation parameters for comparison data. ... 17

Table S2: Prior distribution parameters setting of $BiTSC^2$ for simulations. ... 18

Table S3: MCMC sampling parameters setting of $BiTSC^2$ for simulations. ... 18

Table S4: Prior distribution parameters setting of $BiTSC^2$ for metastatic colorectal cancer dataset. ... 19

Table S5: MCMC sampling parameters setting of $BiTSC^2$ for metastatic colorectal cancer dataset. 19

**Supplementary References** ... **20**

### Supplementary Methods

#### Supplementary Note 1: Inference of parameters

##### Sampling of subclone assignment $C$

We assume $C_n$, the variable indicating the origin subclone of cell $n$, follow the Categorical prior Distribution with parameter $\phi$, which is a vector with length-$K$ and summation of 1. $\phi$ describes the subclonal prevalence, where each element $\phi_k$ represents the proportion of cells from subclone $k$. Then we introduce an additional parameter, $\theta_k$, for each $\phi_k$, and denote the vector comprised of all $\theta_k$ as $\Theta$. We set each $\theta_k$ from an independent Gamma$(\gamma, 1)$ prior distribution, and make $\phi_k = \theta_k / \sum_{i=1}^{K} \theta_i$. This is equivalent to setting $\phi_k$ to a symmetric Dirichlet$(\gamma, \gamma, \cdots, \gamma)$ prior distribution with mean and variance of $(1/K, 1/K, \cdots, 1/K)$. The purpose of introducing $\Theta$ is that we can update one element of $\Theta$ at a time, but to sample $\phi$ directly, all elements of the vector need to be updated for each sampling due to the restriction of the sum of elements as 1. So we take $\Theta$ instead of $\phi$ to work on (Zeng *et al.*, 2019).

Then each element $C_n$ ($n \in \{1, 2, \cdots, N\}$) of $C$ is taken from the Categorical distribution with parameter $\phi$, and subclonal fraction $\phi$ can be updated by updating the parameter $\Theta$. So we update $\Theta$ before we update $C$. We update the one element $\theta_k$ of $\Theta$ at a time. Since the full conditional samples of $\theta_k$, i.e., $p(\theta_k | D, X, C, \Psi, \Omega_{-\theta_k})$ can not be directly sampled, we take Metropolis-Hastings sampling method for sampling $\Theta$.

For the Metropolis-Hastings sampling of $\Theta$, based on the current sample $\theta_k$ in Markov chain, a new $\theta_k^*$ is proposed from the transition function $f(\theta_k^* | \theta_k, \lambda)$, where $f(\theta_k^* | \theta_k, \lambda)$ is the density function of distribution Gamma$(\lambda\theta_k, 1/\lambda)$ with center $\theta_k$ and variance $\theta_k/\lambda$. The tuning parameter $\lambda$ controls the proposing step size, and a larger $\lambda$ value usually leads to a higher acceptance rate. In our implementation, we adaptively adjust its value to keep the acceptance rate in a reasonable range to ensure effective mixing of Markov chains (Zeng *et al.*, 2019).

After updating and sampling each value of $\Theta$, since the sampling space for subclone assignment is discrete and very small, we perform Gibbs sampling on $C$ by calculating the probability of the subclone that each cell $n$ may belong to. Since the cells are independent, we update one element in $C$ each time. For each cell $n$, we calculate $p(C_n = k | D, X, \Psi, \Omega_{-C_n})$ for all possible $k$ ($k \in \{1, 2, \cdots, K\}$) and use them as weights to sample a new $C_n$.

##### Sampling of SNV and CNA origin matrices $L^o$, $Z^o$

Since the sampling spaces of combinations of $(k, v)$ for CNA and SNV status are also discrete and relatively small, then we perform discrete Gibbs sampling by calculating the probability of each possible status.

Because the loci are independent, we can sample and update $Z^o$ row by row when sampling the status of SNV. For each locus $m$, we calculate the posterior probability $p(Z_m^o = (k, v)|D, X, \Psi, \Omega_{-Z_m^o})$ for each state combination $(k, v)$, and use these posterior probability values as weights to sample a new $Z^o$. The sampling process of $L^o$ is similar to that of $Z^o$, but it samples and updates all the loci in a segment together instead of each row at a time.

For the hyper-parameter $\pi$ of $L^o$, we apply Gibbs sampling to update because we can write the fully conditional distribution in the form of Beta distribution as follows:

$$p(\pi|L^o) \sim \text{Beta}(u + \alpha, S - u + \beta),$$

where $S$ is the number of genome segments, and $u$ is the number of segments without CNA (Zeng *et al.*, 2019).

##### Sampling of clone tree $\mathcal{T}$

Since the sampling space of phylogenetic tree is also discrete, but the size increases rapidly with the growth of the number of subclones $K$, it will be a huge computational burden explore all the discrete values of the tree and calculate the posterior probabilities. Here, we adopt a mixed sampling method for the tree, randomly selecting Metropolis-Hastings sampling and slice sampling.

For Metropolis-Hastings sampling, we adopt the following sampling method: randomly select a leaf node and reconnect it to a randomly selected parent node. However, Metropolis-Hastings sampling may fall into a local tree structure, so we adopt the following slice sampling method as another sampling method:

(1) randomly generate a parameter $\sigma \sim \text{Uniform}(0,\ p(\mathcal{T}|D, X, \Psi, \Omega_{-\mathcal{T}}))$,

(2) randomly and repeatedly sample a $\mathcal{T}^*$ from tree space and accept $\mathcal{T}^*$ if $p(\mathcal{T}^*|D, X, \Psi, \Omega_{-\mathcal{T}}) \geq \sigma$.

Slice sampling enables our sampler to make a bigger jumps to avoid falling into local mode. According to empirical analysis, the combination of Metropolis-Hastings sampling and slice sampling not only increases the sampler's mobility, but also produces a higher acceptance rate and is robust in a variety of simulation situations.

##### Sampling of dropout rate $\rho$

Since the full conditional distribution of $\rho$ is difficult to sample directly, we use Metropolis sampling to update $\rho$. Assuming that $\rho_0$ is the sample of the current estimated dropout rate in MCMC chain, we randomly and uniformly sample a new sample $\rho_1$ in the interval $[0, 1]$, calculate the ratio of the posterior probability of $\rho$ (i.e., $p(\rho|D, X, \Psi, \Omega_{-\rho})$) before and after sampling to judge whether to accept the new sample $\rho_1$ according to the Metropolis sampling criterion.

#### Supplementary Note 2: Heuristic initialization process for MCMC parallel chains

We use heuristic initialization for each parallel chain before MCMC sampling. We calculated the VRF at each locus in each cell based on the total reads and mutation reads. Generally, cells from the same subclone have similar variant reads frequency (VRF). Therefore, based on the observed VRF matrix, we use Gaussian mixture model to cluster the cells and obtain the subclone cell assignment to initialize $C$. Then we obtain the VRF of each subclone at each locus, and use the minimum spanning tree (MST) algorithm to construct the subclonal evolutionary path to initialize the clone tree $\mathcal{T}$. After initializing $C$ and $\mathcal{T}$ for each chain, CNA and SNV are randomly allocated on the tree, and then MCMC sampling optimization is performed. We use this heuristic optimization method to divide the more similar cells into the same subclone first, instead of randomly assigning cells, so as to avoid optimizing different subclone genotypes by cells from the same subclone resulting in different subclones sharing the same genotype, which makes the optimization result more accurate.

#### Supplementary Note 3: Derivation of the fully conditional distribution for all model parameters

(1) $\pi$:

the posterior distribution of $\pi$ is (with prior $\text{Beta}(\alpha, \beta)$):

$$\begin{aligned} p(\pi|L^o) &\propto p(L^o|\pi)p(\pi) \\ &\propto \pi^u(1-\pi)^{S-u}p(\pi) \\ &\propto \text{Beta}(u+\alpha, S-u+\beta), \end{aligned}$$

where $S$ is the number of genome segments, and $u$ is the number of segments without CNA.

(2) $\Theta$:

the posterior distribution of $\Theta$ is:

$$\begin{aligned} p(\Theta|D, X, \Psi, C, \Omega_{-\Theta}) &= p(\Theta|C) \\ &\propto p(C|\Theta)p(\Theta) \\ &\propto p(\Theta)\prod_k (\frac{\theta_k}{\sum_k \theta_k})^{N_k}, \end{aligned}$$

where $N_k$ is the number of cells belong to subclone $k$.

(3) $Z^o$:

we update $Z^o$ row by row:

$$\begin{aligned} p(Z^o_m|D,X,\Psi,C,\Omega_{-Z^o_m}) &= p(Z^o_m|D,X,\Psi,C,\mathcal{T}) \\ &\propto p(Z^o_m)\prod_n p(x_{mn}|d_{mn},p_{mn}) \end{aligned}$$

(3) $L^o$:

we update rows of $L^o$ in the same segment together. Consider the loci in segment $\Delta_i$ and assume they share the same CNA status $L^o_{\Delta_i}$:

$$\begin{aligned} p(L^o_{\Delta_i}|D,X,\Psi,C,\Omega_{-L^o_{\{m:m\in\Delta_i\}}}) &= p(L^o_{\Delta_i}|D,X,\Psi,C,\mathcal{T},\rho) \\ &\propto p(L^o_{\Delta_i})\prod_{m\in\Delta_i}\prod_n p(x_{mn}|d_{mn},Z^o_m,L^o_m=L^o_{\Delta_i},C,\mathcal{T}) \\ &\times \prod_{m\in\Delta_i}\prod_n p(d_{mn}|\psi_n,L^o_m=L^o_{\Delta_i},C,\mathcal{T},\rho). \end{aligned}$$

(4) $\mathcal{T}$:

the posterior distribution of $\mathcal{T}$ is:

$$\begin{aligned} p(\mathcal{T}|D,X,\Psi,C,\Omega_{-\mathcal{T}}) &= p(\mathcal{T}|D,X,\Psi,C,L^o,Z^o,\rho) \\ &\propto p(\mathcal{T})\prod_{m,n} p(x_{mn}|d_{mn},C,L^o,Z^o,\mathcal{T})p(d_{mn}|\psi_n,C,L^o,\mathcal{T},\rho) \end{aligned}$$

(5) $\rho$:

the posterior distribution of $\rho$ (with prior uniform on [0,1]) is:

$$\begin{aligned} p(\rho|D,X,\Psi,C,\Omega_{-\rho}) &= p(\rho|D,X,\Psi,C,L^o,Z^o) \\ &\propto p(\rho)p(d_{mn}|\psi_n,C,L^o,\mathcal{T},\rho) \end{aligned}$$

#### Supplementary Note 4: Model selection

We have collected MCMC samples with different number of subclones , but we need to solve the model selection problem. If only the maximum likelihood of the model is considered, it will lead to overfitting issue, because the model with the more subclones are more likely to yield an improved likelihood. There are other selection criteria proposed by researchers, such as maximum posterior posteriori (MAP) (Marass *et al.*, 2016). However, MAP approach relies on the selection of prior distribution to a large extent (Zeng *et al.*, 2019). Schwarz *et al.* (1978) proposed a criterion, called

Bayesian Information Criterion (BIC), to select models by a large sample approximation of the respective marginal distribution of the data $p(x) = E_{h(\theta)}[f(x|\theta)]$. BIC is quite different from that of measures of posterior predictive accuracy, which based on the weight of sample distribution with the prior density. For large sample sizes $n$, there is $\ln p(x) \simeq -2\ln f(x|\hat{\theta}) + p\ln n$, where $\hat{\theta}$ is the maximum likelihood estimate of $\theta$ and $p$ is the number of parameters in $\theta$. Then the definition of BIC can be

$$BIC = -2\ln f(x|\hat{\theta}) + p\ln n.$$

In this study, we adopt modified version of BIC proposed by Carlin and Louis (2009), using the posterior mean of loglikehood instead of maximum for avoiding simulation-based maximization to perform model selection, which is defined as (Turkman *et al.*, 2019)

$$BIC = -2E_{\theta|x}[\ln f(x|\theta)] + p\ln n.$$

BIC not only prefers the model with large loglikehood values, but also adds the penalty on the number of free model parameters to punish the complexity of the model and shows heavily penalization on more complicated models for large samples. Then according to the definition of BIC, we choose model with small value of BIC.

#### Supplementary Note 5: Evaluations of ARI

For comparison of the accuracy of subclone assignment, we used ARI (Rand, 1971; Qiu *et al.*, 2017) to measure the similarity between ground truth and estimation of $C$. Assume there are two partitions, $P^{(1)} = \{P_1^{(1)}, P_2^{(1)}, \cdots, P_r^{(1)}\}$ and $P^{(2)} = \{P_1^{(2)}, P_2^{(2)}, \cdots, P_s^{(2)}\}$ that divide set A into $r$ and $s$ groups, respectively. Let $n_{ij}$ represents the overlap number in $P_i^{(1)}$ and $P_j^{(2)}$, that is, $n_{ij} = |P_i^{(1)} \cap P_j^{(2)}|$, then the set of $\{n_{ij}|i \in \{1, \cdots, r\}, j \in \{1, \cdots, s\}\}$ describes overlap between $P^{(1)}$ and $P^{(2)}$. We then define the number of cells within group $i$ from partition $P^{(1)}$, as $a_i = \sum_{j=1}^{s} n_{ij}$, and the number of cells within subclone $j$ from partition $P^{(2)}$, as $b_j = \sum_{i=1}^{r} n_{ij}$. Then ARI of the two partition can calculated by

$$\text{ARI}(P^{(1)}, P^{(2)}) = \frac{\sum_{ij}\binom{n_{ij}}{2} - [\sum_i \binom{a_i}{2} \sum_j \binom{b_j}{2}]/\binom{n}{2}}{\frac{1}{2}[\sum_i \binom{a_i}{2} + \sum_j \binom{b_j}{2}] - [\sum_i \binom{a_i}{2} \sum_j \binom{b_j}{2}]/\binom{n}{2}}.$$

The value of ARI is in the region of [0,1]. A larger value indicates better assignment.

#### Supplementary Figures

**Figure S1: The ground truth of simulation datasets in G1-G3, containing subclonal phylogenetic tree and genotype matrix of CNA $L$ and SNV $Z$.**

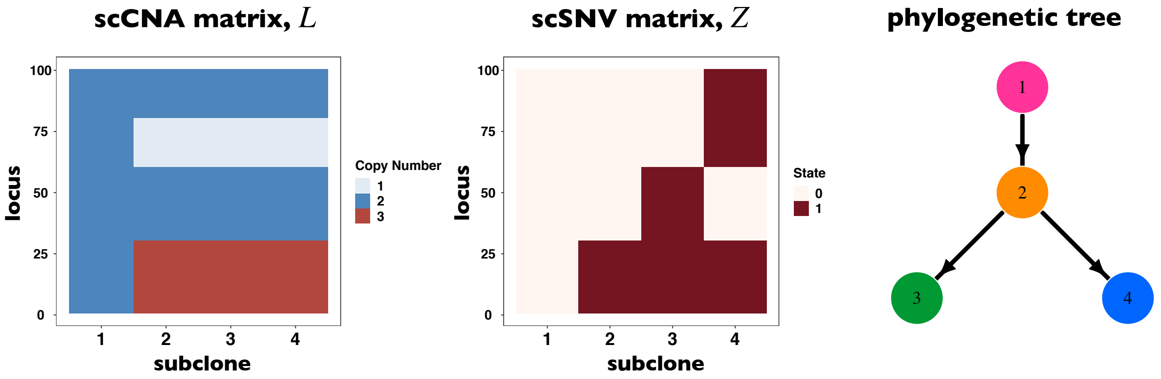


**Fig. S1.** The ground truth of simulation datasets in G1-G3, containing subclonal phylogenetic tree and genotype matrix of CNA $L$ and SNV $Z$.

**Figure S2: The ground truth of simulation datasets in G4, containing subclonal phylogenetic tree and genotype matrix of CNA $L$ and SNV $Z$.**

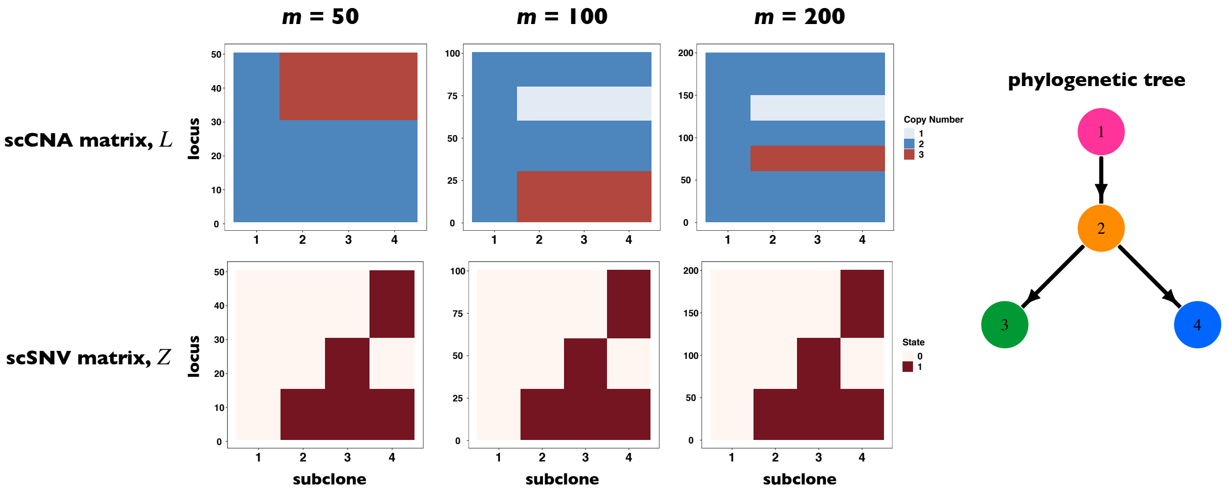


**Fig. S2.** The ground truth of simulation datasets in G4, containing subclonal phylogenetic tree and genotype matrix of CNA $L$ and SNV $Z$.

**Figure S3: The ground truth of simulation datasets in G5, containing subclonal phylogenetic tree and genotype matrix of CNA $L$ and SNV $Z$.**

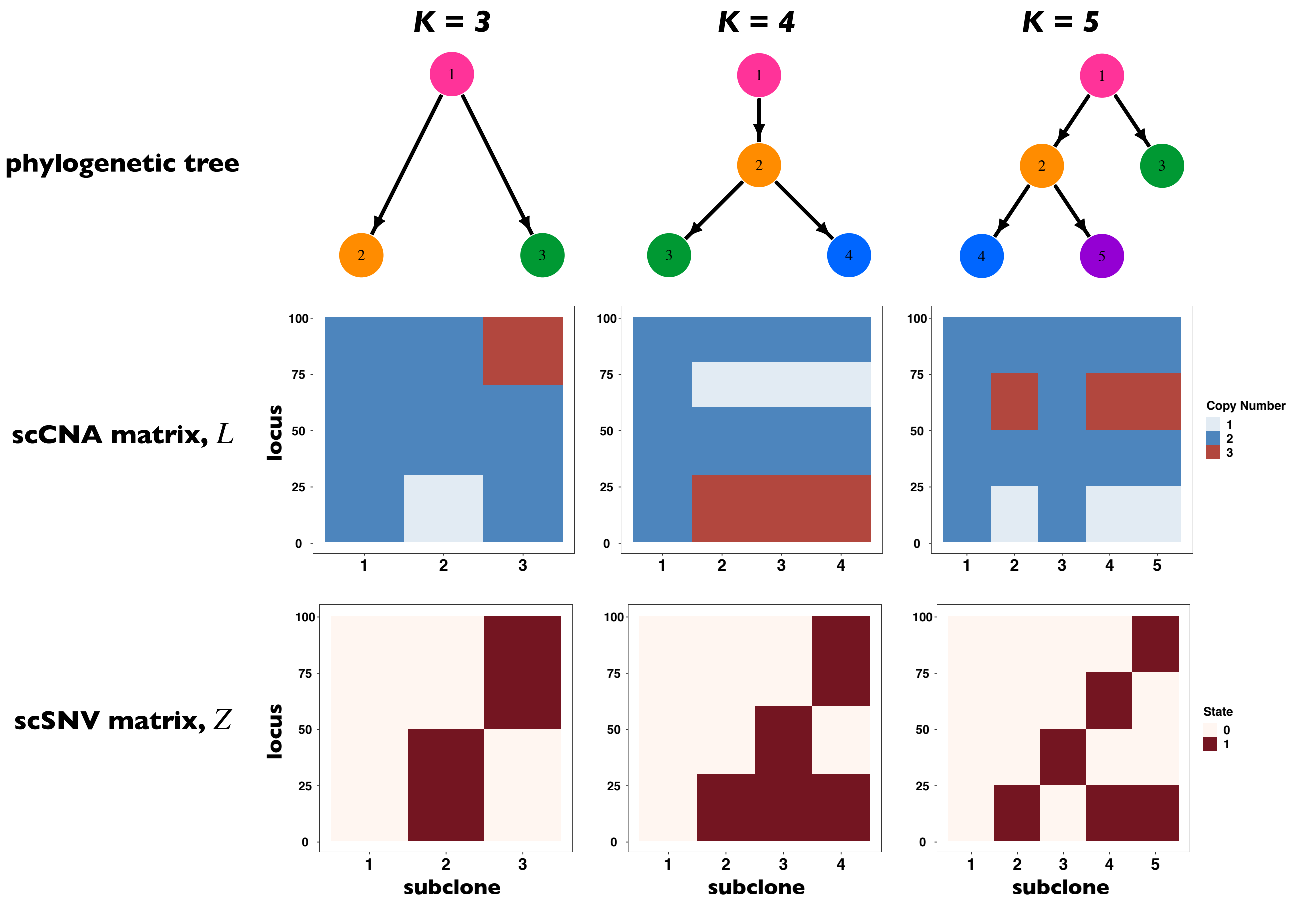


**Fig. S3.** The ground truth of simulation datasets in G5, containing subclonal phylogenetic tree and genotype matrix of CNA $L$ and SNV $Z$.

**Figure S4: The ground truth of simulation datasets in G6, containing subclonal phylogenetic tree and genotype matrix of CNA $L$ and SNV $Z$.**

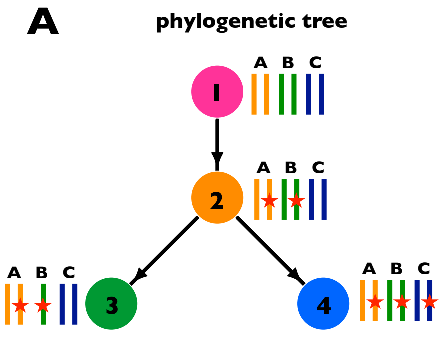


**Fig. S4.** The ground truth of simulation datasets in G6, containing phylogenetic tree and subclonal genotype matixes of CNA and SNV.

**Figure S5: Comparison of performs on G1-G5 for subclone assignment and scSNV genotype recovery among *BiTSC*$^2$ with updating one locus at a time, RobustClone and BEAM.**

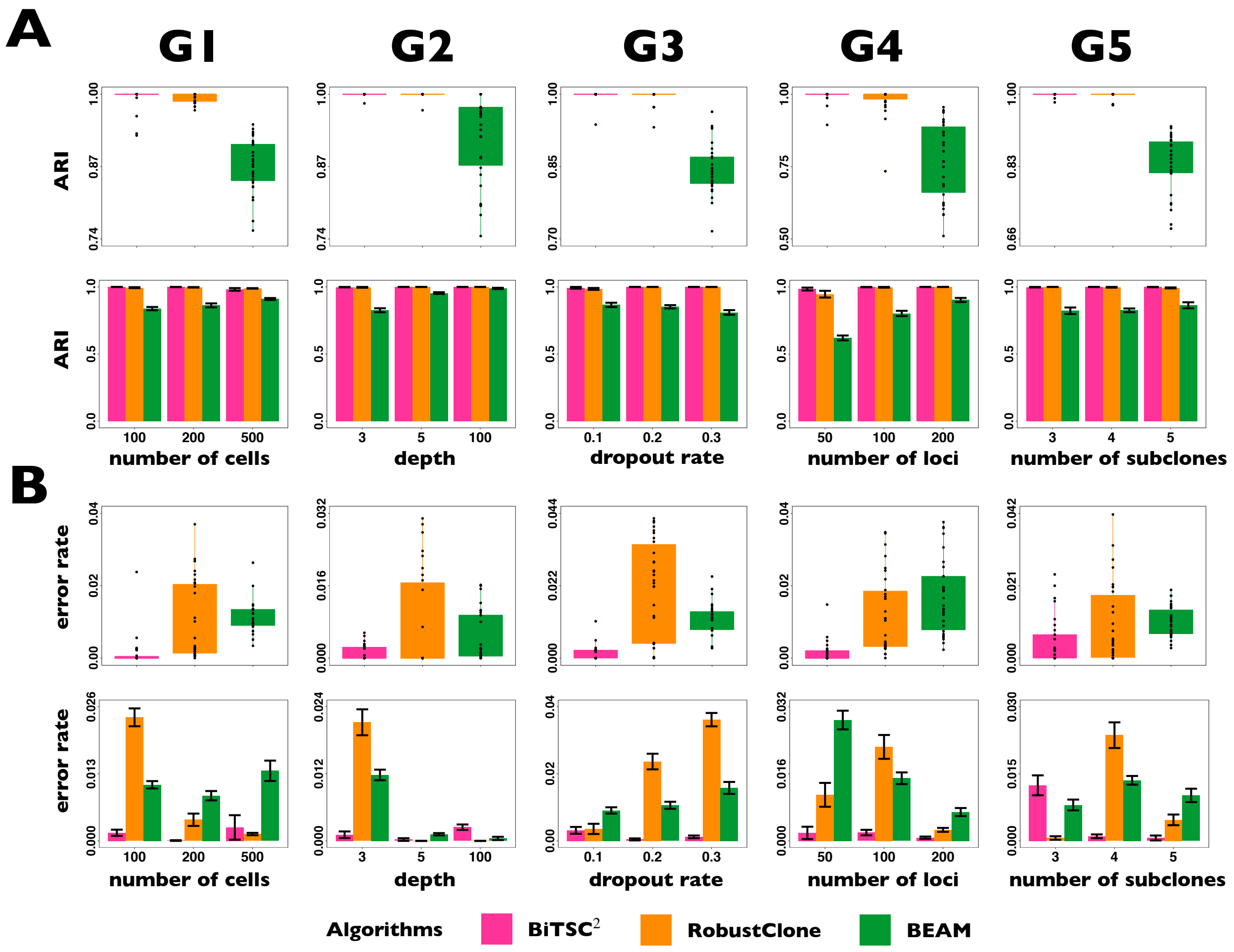


**Fig. S5.** Comparison of performs on G1-G5 for subclone assignment and scSNV genotype recovery among *BiTSC*$^2$ with updating $L^o$ one locus at a time, RobustClone and BEAM. **(A)** The boxplot and barplot of three algorithms for ARI of subclone assignment. **(B)** The boxplot and barplot of three algorithms for error rate of recovered scSNV genotype matrix.

**Figure S6: Comparison of performs on G6 for subclone assignment and scSNV genotype recovery among *BiTSC*$^2$ with updating one locus at a time, RobustClone and BEAM.**

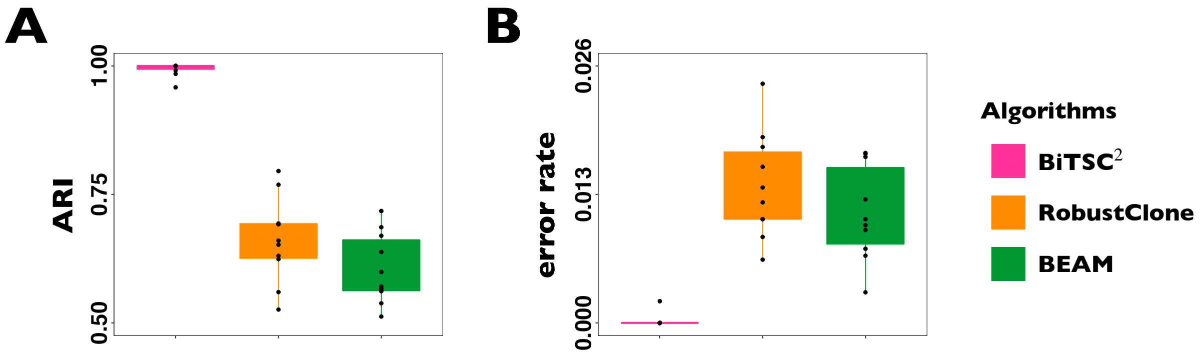


**Fig. S6.** Comparison of performs on G6 for subclone assignment and scSNV genotype recovery among $BiTSC^2$ with updating $L^o$ one locus at a time, RobustClone and BEAM. **(A)** The boxplot of three algorithms for ARI of subclone assignment. **(B)** The boxplot of three algorithms for error rate of recovered scSNV genotype matrix.

**Figure S7: The BIC of metastatic colorectal cancer data calculated in Model selection step.**

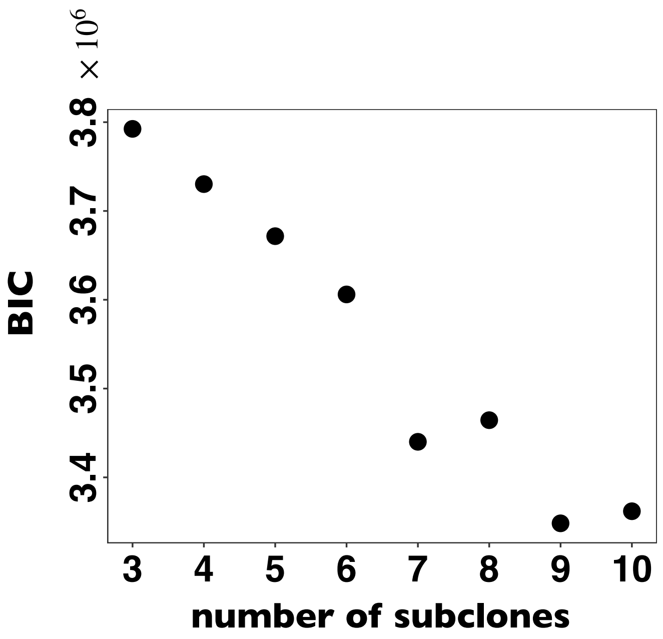


**Fig. S7.** The BIC of metastatic colorectal cancer data calculated in Model selection step.

**Figure S8: The phylogeny tree inferred by SCITE and SCARLET on the dataset from a Metastatic Colorectal Cancer Patient.**

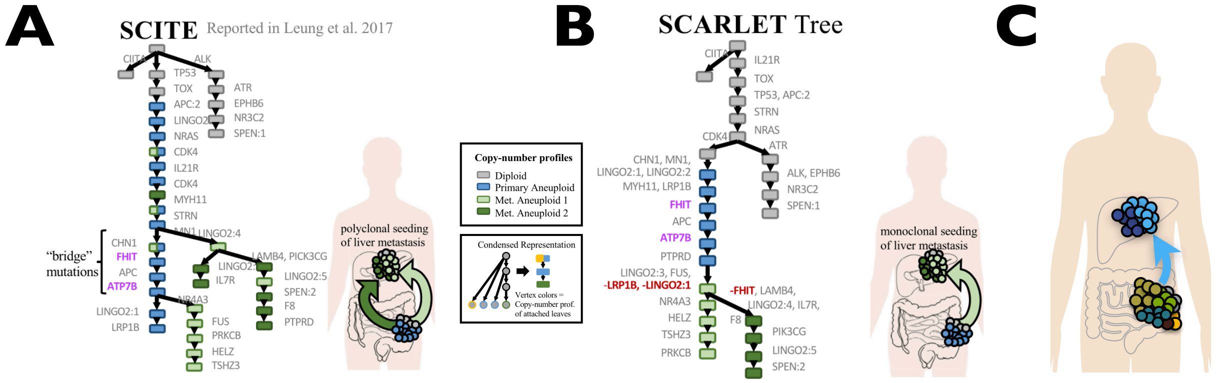


**Fig. S8.** The phylogeny tree inferred by SCITE and SCARLET on the dataset from a Metastatic Colorectal Cancer Patient. **(A)**The phylogeny tree inferred by SCITE in (Leung *et al.*, 2017), where two distinct branches of metastatic cells suggest polyclonal seeding of liver metastasis. **(B)** The phylogeny tree inferred by SCARLET with all metastatic cells contained in a single branch, which suggests monoclonal seeding of the liver metastasis. **(C)** The phylogeny tree inferred by $BiTSC^2$ in Figure 2A suggests monoclonal seeding of the liver metastasis. Figure AB are adapted from Satas *et al.* (2020).

**Figure S9: The dropout rate $\rho$ estimated by $BiTSC^2$ on the datasets from G3 groups.**

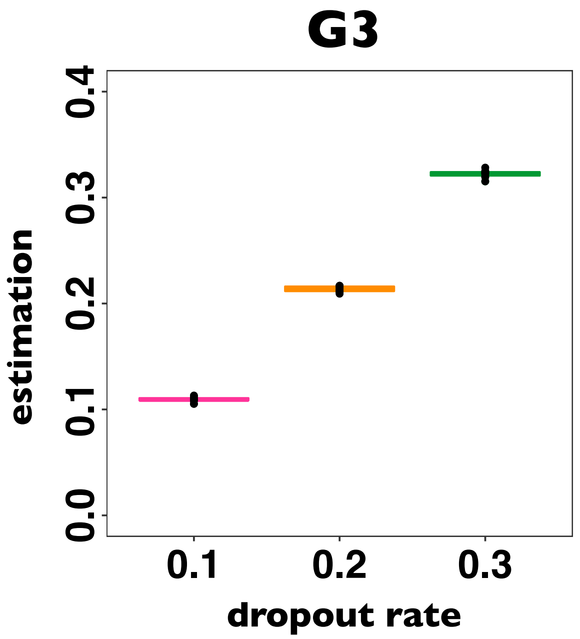


**Fig. S9.** The dropout rate $\rho$ estimated by $BiTSC^2$ on the datasets from G3 groups.

#### Supplementary Tables

##### Table S1: The setting of simulation parameters for comparison data.

Table S1: The setting of simulation parameters for comparison data.

| | change factor | | | control factors | | | |
|---|---|---|---|---|---|---|---|
| G1 | $n$ | | | $\psi$ | $\rho$ | $m$ | $K$ |
| | 100 | 200 | 500 | 3 | 0.2 | 100 | 4 |
| G2 | $\psi$ | | | $n$ | $\rho$ | $m$ | $K$ |
| | 3 | 5 | 100 | 100 | 0.2 | 100 | 4 |
| G3 | $\rho$ | | | $n$ | $\psi$ | $m$ | $K$ |
| | 0.1 | 0.2 | 0.3 | 100 | 3 | 100 | 4 |
| G4 | $m$ | | | $n$ | $\psi$ | $\rho$ | $K$ |
| | 50 | 100 | 200 | 100 | 3 | 0.2 | 4 |
| G5 | $K$ | | | $n$ | $\psi$ | $\rho$ | $m$ |
| | 3 | 4 | 5 | 100 | 3 | 0.2 | 100 |

#### Table S2: Prior distribution parameters setting of $\boldsymbol{BiTSC^2}$ for simulations.

Table S2: Prior distribution parameters setting of $BiTSC^2$ for simulations.

| Parameters | Value |
|---|---|
| Maximum possible mutant copies ($M_s$) | 2 |
| Maximum possible total copies ($M_c$) | 4 |
| Dirichlet prior parameter of $\phi$ ($\gamma$) | 1.5 |
| Beta prior parameter of $\pi$ ($\alpha, \beta$) | (10000,1) |
| prior parameter of $Z^o$ ($\zeta$) | 0.01 |

#### Table S3: MCMC sampling parameters setting of $\boldsymbol{BiTSC^2}$ for simulations.

Table S3: MCMC sampling parameters setting of $BiTSC^2$ for simulations.

| Parameters | Value |
|---|---|
| Number of chains | 5 |
| Temperature increment ($\Delta T$) | 0.35 |
| Sample size for posterior inference | 500 |
| Burn-in sample size | 500 |
| Sample size for tuning adaptive parameter | 500 |
| Interval to perform chain swap | 30 |
| Probability of tree slice sampling | 0.15 |
| Probability of tree Metropolis-Hastings sampling | 0.85 |

#### Table S4: Prior distribution parameters setting of $BiTSC^2$ for metastatic colorectal cancer dataset.

Table S4: Prior distribution parameters setting of $BiTSC^2$ for metastatic colorectal cancer dataset.

| Parameters | Value |
|---|---|
| Maximum possible mutant copies ($M_s$) | 5 |
| Maximum possible total copies ($M_c$) | 10 |
| Dirichlet prior parameter of $\phi$ ($\gamma$) | 1.5 |
| Beta prior parameter of $\pi$ ($\alpha, \beta$) | (10000,1) |
| prior parameter of $Z^o$ ($\zeta$) | 0.01 |

#### Table S5: MCMC sampling parameters setting of $BiTSC^2$ for metastatic colorectal cancer dataset.

Table S5: MCMC sampling parameters setting of $BiTSC^2$ for metastatic colorectal cancer dataset.

| Parameters | Value |
|---|---|
| Number of chains | 5 |
| Temperature increment ($\Delta T$) | 0.35 |
| Sample size for posterior inference | 500 |
| Burn-in sample size | 500 |
| Sample size for tuning adaptive parameter | 500 |
| Interval to perform chain swap | 30 |
| Probability of tree slice sampling | 0.15 |
| Probability of tree Metropolis-Hastings sampling | 0.85 |
